# *Rhizoctonia theobromae* associated with a severe witches’ broom outbreak in cassava of the Brazilian rainforest and evidence of its Southeast Asian origin

**DOI:** 10.1101/2025.02.14.638269

**Authors:** Saulo Alves Santos de Oliveira, Samar Sheat, Paolo Margaria, Adilson Lopes Lima, Jackson de Araújo dos Santos, Hermínio Souza Rocha, Helton Fleck da Silveira, Cristiane Ramos de Jesus, Stephan Winter

## Abstract

Cassava witches’ broom disease (CWBD) has emerged as a significant threat to cassava production in the Oiapoque region of Amapá, Brazil. Diseased plants exhibit stunted growth, vascular necrosis, abnormal shoot proliferation, and distinctive broom-like appearance. This study aimed to characterize the disease and identify its causal agent(s). The assessment by high-throughput sequencing of total nucleic acids (DNA and RNA) from affected cassava tissues revealed *Rhizoctonia theobromae* (syn. *Ceratobasidium theobromae*) associated with CWBD. PCR-based detection using species-specific primers confirmed the presence of *R. theobromae* in 74% of symptomatic samples. Genetic diversity analysis based on the Ca2+/calmodulin-dependent protein kinase gene showed low variability among Brazilian isolates compared with Asian populations, suggesting their recent introduction. This first report of *R. theobromae* causing CWBD in Brazil follows a recent report of the disease from French Guiana and highlights the urgent need for effective measures to prevent the further spread of this emerging pathogen, which poses a deadly threat to cassava cultivation in the rainforest, a significant risk to cassava production in Brazil, and regions with similar eco-climatic conditions.

## INTRODUCTION

*Manihot esculenta* Crantz, commonly known as cassava, serves as a primary food source for approximately one billion people across 105 nations in the tropics. Following rice and maize, it ranks as the third most significant carbohydrate provider (Bayata, 2019) and has substantial economic and social value, particularly in developing nations (Muiruri et al. 2021). Despite being primarily cultivated as a staple food, in recent years, it has been increasingly utilized as a source of bioenergy, starch, alcohol, biopolymers, cosmetics, and beyond (Fathima et al. 2023). Brazil is the sixth largest producer of cassava tuberous roots globally, with a production of 17.82 million tons (FAO 2022), which is almost exclusively produced to satisfy domestic market demands. Thus, cassava is a vital source of food and income for the Brazilian population.

This hitherto unknown disease is affecting cassava in the Indigenous lands of the Oiapoque region in Brazil bordering French Guiana, where it presents a serious threat to the livelihoods of people relying on cassava as their main diet. This was expressed at the XXIX Assembly for the Evaluation and Planning of Indigenous Peoples and Organizations in the municipality of Oiapoque, Amapá, Brazil, on March 11 and 12, 2023, and an urgent request was made to investigate the origin of the disease and provide solutions to mitigate its impact on the population. Cassava grown in the forest exhibited abnormal growth, with deformed leaves on the shoots. Flushing leaves on short internodes form clusters and appear as broom-like bundles, giving rise to the name cassava witches’ broom disease (CWBD). Leaf yellowing and vascular necrosis are additional symptoms of this disease. Previously, similar symptoms have been observed in cassava grown in Southeast Asia (SEA) (Pardo et al. 2023) and were recently reported in French Guiana (Pardo et al. 2024). Eventually, the disease leads to a decline and wilting of leaves and stems, resulting in very thin and small roots that compromise the yield and quality of the harvest.

A cassava witches’ broom-like disease was reported in the 1940s in Brazil (Silberschmidt and Campos 1944), and later, phytoplasma (16SrIII-B group) organisms were associated with the characteristic “oversprouting symptoms” (Flores et al. 2013). In contrast to the earlier disease report, the new cassava disease in Oiapoque comprises vascular necrosis and wilting, leading to decay and eventual plant death. Cassava affected by CWBD in SEA, including in Cambodia, Laos, Vietnam, Thailand, and the Philippines, was assumed to be a phytoplasma disease (Pardo et al. 2023); however, further biological evidence is needed. Recently, a significant breakthrough in clarifying the root cause of CWBD in ‘SE’ Asia was reported. Using a combination of high-throughput sequencing (HTS), isolation, and culture techniques, Leiva et al. (2023) and Gil-Ordóñez et al. (2024) provided compelling evidence that the fungus *Rhizoctonia theobromae* (syn = *Ceratobasidium theobromae*) was the cause of CWBD in SEA. High-resolution microscopy has revealed that the fungus occupies xylem cells, directly correlating with the observed vascular necrosis (Gil-Ordóñez et al. 2024). Further proof was provided by transmitting *Rhizoctonia/Ceratobasidium* through wedge grafting to infect plants (Leiva et al. 2023). Landicho et al. (2024) studied CWBD in the Philippines using PCR and HTS and showed that the microbial composition of CWBD-affected cassava leaves was dominated by *Rhizoctonia/Ceratobasidium*, thus confirming the findings from Southeast Asia.

Our research on CWBD in cassava from Oiapoque was guided by these earlier findings. To gain a comprehensive understanding of the pathobiome of cassava from this region, we employed an HTS approach for a global and unbiased assessment of the cassava metagenome. A substantial portion of the DNA sequences not mapped to the cassava genome was from *Rhizoctonia theobromae* (syn = *Ceratobasidium theobromae*), whereas DNA from other microorganisms was also found in the sample. We further employed a genetic approach to trace the origin of this quarantine pathogen using Ca2 +/calmodulin-dependent protein kinase (CAMK/CAMKL) sequences to provide critical insights into disease etiology, source of origin, and potential impact on cassava production in Brazil.

## MATERIALS AND METHODS

Fifty samples were collected from the villages of Kuai Kuai, Ariramba, Galibi, Lençol, Ahumã, Anawerá, and Yanawaká in the Oiapoque Region of Amapá (Figure 1).

**Figure 1.**
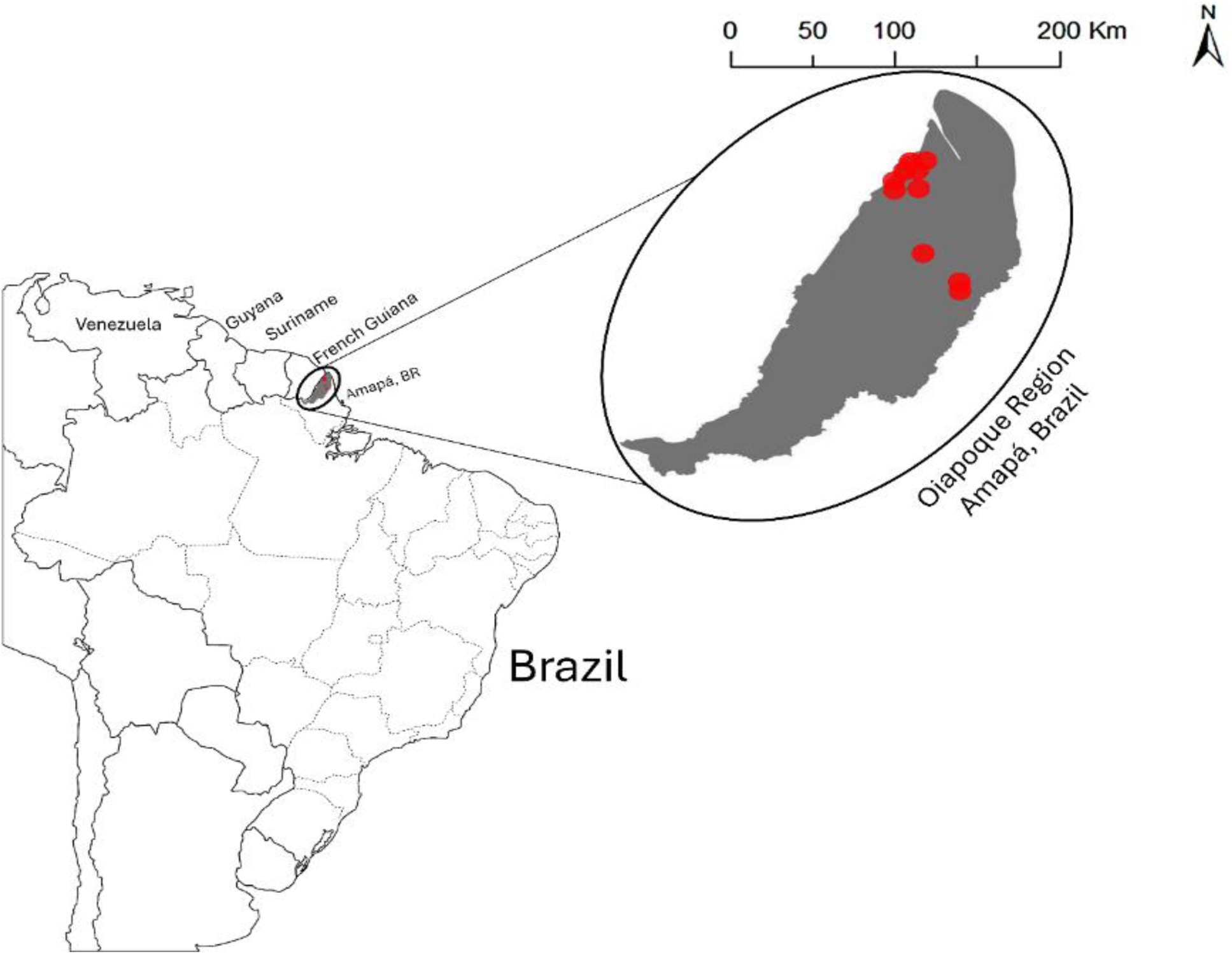
Cassava samples were collected from diseased plants in Kuai Kuai, Ariramba, Galibi, Lençol, Ahumã, Anawerá, and Yanawaká. The red dots indicate the collection sites.

Leaves, petioles, and stems from symptomatic cassava plants were collected from the fields (Table 1), and after registration in SISGEN (**AFB8348**), packaged in sealed containers and transported to Embrapa Mandioca e Fruticultura in Cruz das Almas-Bahia, Brazil.

**Table 1.**
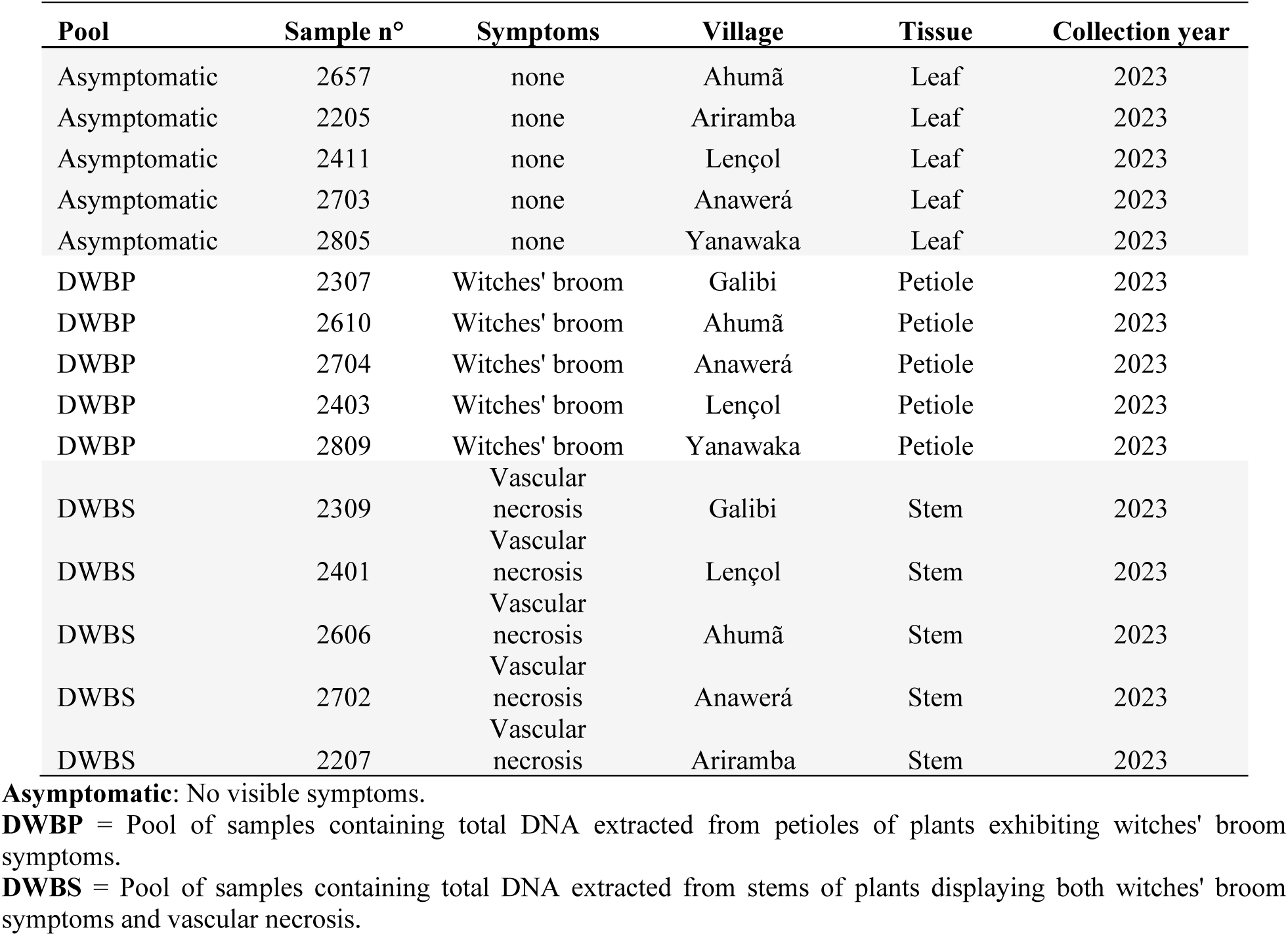
Overview of the samples collected from different villages in 2023.

The materials were flash frozen in liquid nitrogen and lyophilized for 72 h at a constant pressure and temperature (54 µHg and -53° C, respectively). Freeze-dried materials were stored at -80 °C prior to shipment to the Leibniz Institute (DSMZ, Germany) for further analysis.

### Pathogen detection based on High throughput sequencing

HTS sequencing and subsequent bioinformatics analysis were conducted at the Plant Virus Department (Leibniz Institute DSMZ, Braunschweig, Germany) using specific workflows developed for pathogen discovery. Total DNA was extracted from each sample (150 mg of stem, leaf, or petiole) using NucleoSpin® Plant II (MACHEREY-NAGEL GmbH & Co. KG, Germany) kit, following the protocol for isolation of “Genomic DNA from plant” provided by the manufacturer. The final total DNA was eluted in 100 µL of elution buffer, checked on a 1% agarose gel (Promega, Germany), quantified using a NanoDrop^TM^ and Qubit^TM^ spectrophotometer (Thermo Fisher Scientific, Waltham, MA, USA), and stored at −20 °C.

HTS of the total DNA from cassava was performed using sample pools composed of similar tissues (leaves, petioles, and stems). For Illumina and Nanopore sequencing, similar amounts of DNA were combined into pools (Table 1) of DNA from non-symptomatic leaves, petioles from witches’ broom plants, and stems with vascular necrosis. Library preparation was performed using the Illumina DNA Prep Kit (FlexM) along with IDT® for Illumina® DNA/RNA UD Indexes Set D. The double-stranded cDNA (ds-cDNA) used for library preparation was generated as following: First, random cDNA was synthesized using the Qiaseq FastSelect -rRNA Plant Kit (Qiagen), RiboLock RNase Inhibitor (Thermo Scientific), Maxima H Minus Reverse Transcriptase (Thermo Scientific), dNTPs (Invitrogen,), an Octamer Primer (Eurofins), and RNase H (Invitrogen). Second-strand synthesis was conducted using the NEBNext Ultra II Non-Directional RNA Second Strand Synthesis Module (NEB), followed by purification using the NucleoSpin Gel and PCR Clean-up Kit (Macherey-Nagel). The generated libraries were quantified using a DNA kit on the Agilent 2100 Bioanalyzer, according to the manufacturer’s instructions.

Sequencing was performed on a NextSeq 2000 instrument (Illumina, San Diego, CA, USA) using paired-end reads (2 × 150 nt with P2 Reagents v3, 300 cycles). Quality control steps included adaptor trimming, error correction, and normalization. Processed reads were then mapped against the *Manihot esculenta* reference genome {(National Center for Biotechnology Information, 2021 – *Manihot esculenta* v8.1, NCBI RefSeq assembly GCF_001659605.2, Annotation Release 101) [cited 2024 Mar 05]. Available from: https://www.ncbi.nlm.nih.gov/datasets/genome/GCF_001659605.2} with unmapped reads retained for *de novo* assembly using Geneious Prime^®^ (v.2024.0; Biomatters).

Nanopore libraries were prepared following protocols provided by Oxford Nanopore Technologies (ONT, UK) and sequenced on the MinION platform. Raw reads were base-called and demultiplexed using Guppy (v6.0.1), followed by quality control with ‘Porechop’ for adaptor trimming, error correction, and normalization. The reads were mapped against the *M. esculenta* reference genome (NCBI RefSeq assembly GCF_001659605.2) using ‘Minimap2’, with unmapped reads retained for further downstream analyses.

### Bioinformatic analysis

To obtain a general overview of the HTS-sequenced samples, the assembled *de novo* contigs generated from Illumina sequencing and processed nanopore reads were analyzed using the DIAMOND BLASTx tool (Buchfink et al., 2015) to identify potential homologs. The resulting BLASTx output was then imported into the MEGAN (MEtaGenome ANalyzer) to obtain a metagenomic overview of species based on sequence hits. MEGAN was used to assign taxonomic identities to the contigs, providing insights into the diversity and composition of the species present in the samples based on sequence data. This combined approach enabled a comprehensive analysis of putative protein-coding sequences and their taxonomic affiliations.

To ensure correct species assignment, trimmed reads and *de novo* assembled contigs were screened against a custom database comprising specific sequences for each potentially pathogenic organism using BLAST.

### Phylogenetic analyses

The rDNA ITS regions (ITS1, 5.8S, and ITS2) of the fungi associated with CWBD-affected cassava were aligned with reference isolates from various species complexes using the MUSCLE algorithm in MEGA v11 (Tamura et al., 2021). Although ITS may not always provide species-level resolution, it is effective in delineating species complexes. Reference sequences from GenBank were selected based on similarity indices to enhance the alignment accuracy. *Scotomyces subviolaceus* and *Heteroacanthella acanthophysa* were included as outgroups to root the phylogenetic analysis. Phylogenetic analysis was conducted using the Maximum Likelihood (ML) method in MEGA v.11 with 1000 bootstrap replicates. The percentage of pairwise identity among the aligned nucleotides was calculated using the substitution model T92+G (Tamura 3-parameter with gamma distribution), selected based on Bayesian Information Criterion (BIC) and Akaike Information Criterion (AIC) scores. The trees were visualized and edited using FigTree v.1.4.4 (available at http://tree.bio.ed.ac.uk/software/figtree).

Partial genome scaffolds obtained from the Illumina and ONT sequence assemblies were compared with those of closely related organisms to achieve a more accurate classification. Pairwise comparisons were conducted between the scaffolds generated from the DWBP and DWBS samples and *Rhizoctonia solani* AG-1 IA (GCF_016906535.1) and *Ceratobasidium theobromae* LAO01 (GCA_037974915.1). Additional identity analysis was performed based on other taxonomically informative genes, such as large subunit ribosomal RNA (LSU), translation elongation factor 1-alpha (*tef1*), RNA polymerase II subunit (*rpb2*), and ATP synthase subunit 6 (*atp6*), to enhance the taxonomic resolution and accuracy of the generated sequences when compared by BLAST with different species and with the reference genomes of *C. theobromae* strains (GCA_037974915.1, GCA_009078325.1, and GCA_012932095.1).

### PCR-based detection

To confirm the presence of the fungus in the collected samples, detection was carried out using the primers CWBD-CIAT-F2 (GGATGAGTTTAATCGCTCTAAC) and CWBD-CIAT-R2 (GCGCTCTGGTGTTTCAAGTTTG) (Leiva et al. 2023), developed based on the amplification of the putative Ca^2+^/calmodulin-dependent protein kinase gene (CAMK/CAMKL) via polymerase chain reaction (PCR). PCR was performed in a total volume of 25 µL, including 2.5 µL of 10x PCR buffer, 0.5 µL of forward primer CWBD-CIAT-F2 (10 µM), 0.5 µL of reverse primer CWBD-CIAT-R2 (10 µM), 1.87 µL of MgCl₂ (50 mM), 0.5 µL of Taq DNA polymerase, 0.5 µL) dNTPs, 16.63 µL, nuclease-free water, and 2.0 µL of template DNA (30 ng/µL).

PCR amplification was conducted with an initial denaturation at 95 °C for 5 min, followed by 35 cycles of denaturation at 95 °C for 1 min, annealing at 52 °C for 1 min, and extension at 72 °C for 1 min. After amplification, a final extension step was performed at 72 °C for 10 min and the reaction was maintained at 12 °C until further processing. The initial denaturation step was adjusted to 95 °C for 5 min to accommodate the use of GoTaq DNA Polymerase. Using this procedure, the presence of *R. theobromae* (*C. theobromae*) in the evaluated samples was verified. The PCR products were purified using the Monarch® PCR & DNA Cleanup kit (NEB), following the manufacturer’s instructions, and sent for ‘Sanger sequencing’ of the PCR products in both directions to a commercial service provider (Microsynth Seqlab GmbH, Germany).

### *Rhizoctonia theobromae* genetic diversity analysis

CAMK/CAMKL sequences obtained from Sanger sequencing were quality checked, assembled, and manually edited to generate a consensus sequence for each isolate. The consensus sequences were compared to those available in GenBank using BLASTn (National Center for Biotechnology Information, 2024) [cited 2024 Oct 02]. Available from: https://www.ncbi.nlm.nih.gov.

The generated consensus sequences were aligned using the MUSCLE algorithm implemented in MEGA v11 alongside isolates responsible for cassava witches’ broom disease (CWBD) in Asia. Sequence ends were manually trimmed based on the shortest aligned sequence to ensure uniform length.

DnaSP ver. 6.12.03 (Rozas et al. 2017) was used to calculate genetic diversity indices. The indices included haplotype (gene) diversity (Hd), calculated as the probability that two randomly chosen haplotypes from a given population were different, and nucleotide diversity (π), calculated as the average number of nucleotide differences per site between two DNA sequences in all the sample pairs in the population.

To compare the relationships between populations from different countries, a haplotype analysis network based on single nucleotide variation (SNV) was built using the median-joining method using PopART 1.7 software (Leigh and Bryant 2015) with zero epsilon as a default parameter. A Nexus file generated by DnaSP was used as an input file for PopART 1.7 software. Coloring of the haplotype network was based on the country of origin of *R. theobromae* CAMK/CAMKL DNA sequences.

## RESULTS

Characteristic symptoms of CWBD were found on cassava from indigenous villages located in the Oiapoque Region, Amapá, Brazil (Figure 2a-c). Symptoms included stunting and proliferation of short, thin, and weak shoots on cassava stems, resulting in the formation of “brooms”, represented by short internodes, aggregated rosette-shaped leaves aggregation and vascular necrosis. In severe cases, a cotton-like mycelium oozed out of the petiole base near the buds (Figure 2 e). The fungus produced characteristic aerial light-yellowish mycelia with hyphal branches at approximately 90° angles (Figure 2 e). These branches exhibited constrictions at their bases and were immediately followed by the septa beyond the point of branching (Figure 2 f). As the disease progresses, chlorosis, wilting and drying of leaves, apical death, and descending plant death occur.

**Figure 2.**
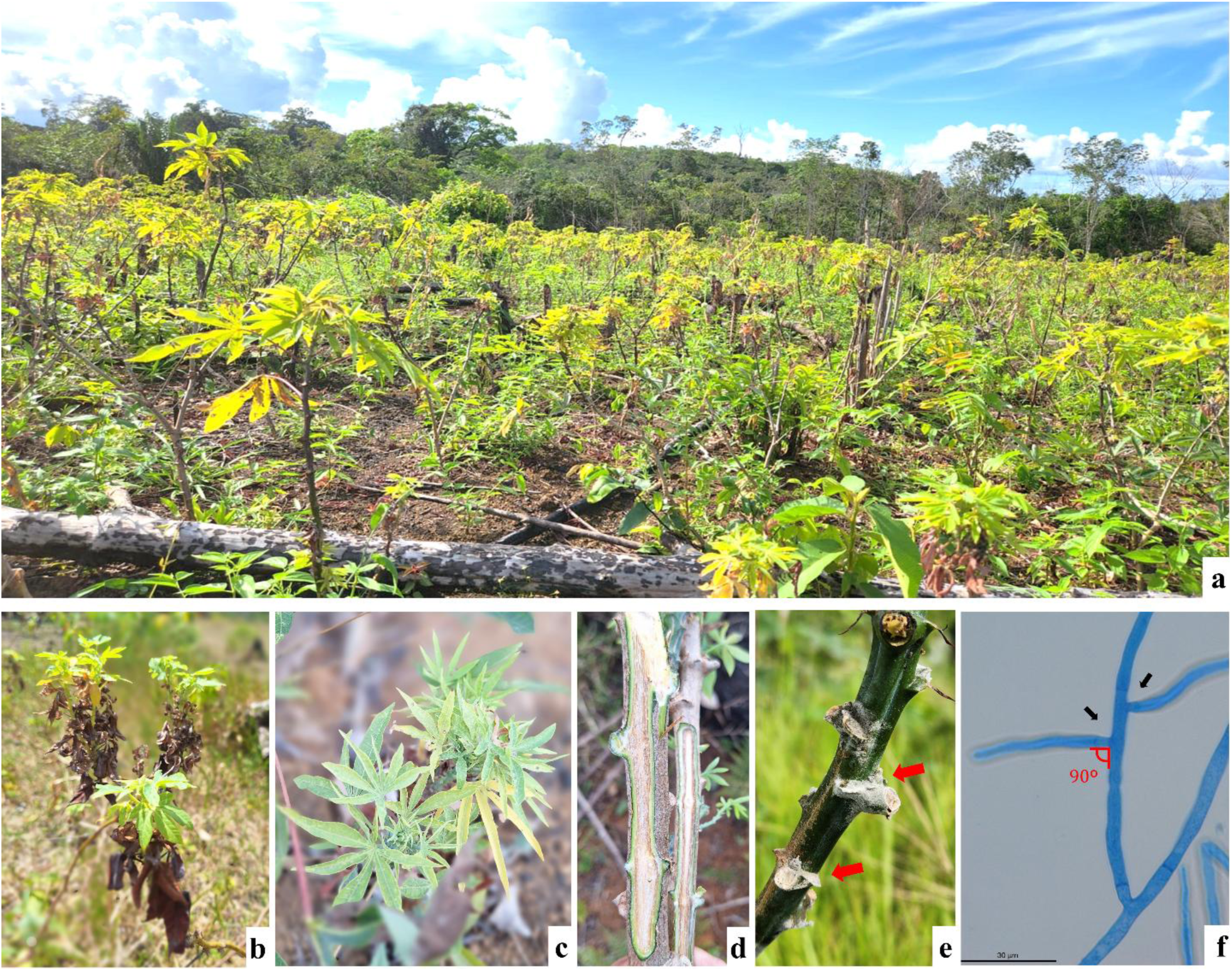
Cassava fields in the indigenous areas of Oiapoque, Amapá, Brazil affected by CWBD. Plants expressing the symptoms of witches’ broom with aggregations of rosette-shaped leaves (**a-c)**; longitudinal section of a cassava stem showing vascular necrosis (**d**); whitish mycelium oozing out of nodes indicating fungal infection (**e) and** typical “*Rhizoctonia*-like” hyphae isolated from petioles, stained with lactophenol blue and observed under light microscope (×400) (**f**).

### CWBD and “oversprouting” disease

This new disease of cassava, CWBD, shares similarities with the "superbrotamento" or “Brazilian witches’ broom”, herein referred to as ‘Oversprouting’. Both result in abnormal shoot proliferation and stunted plants but “Oversprouting” is characterized by generalized stunting, excessive shoot proliferation, and leaves that are chlorotic, deformed, and smaller than normal. The new stems are unusually thin and produced in large numbers (Figure 3).

**Figure 3.**
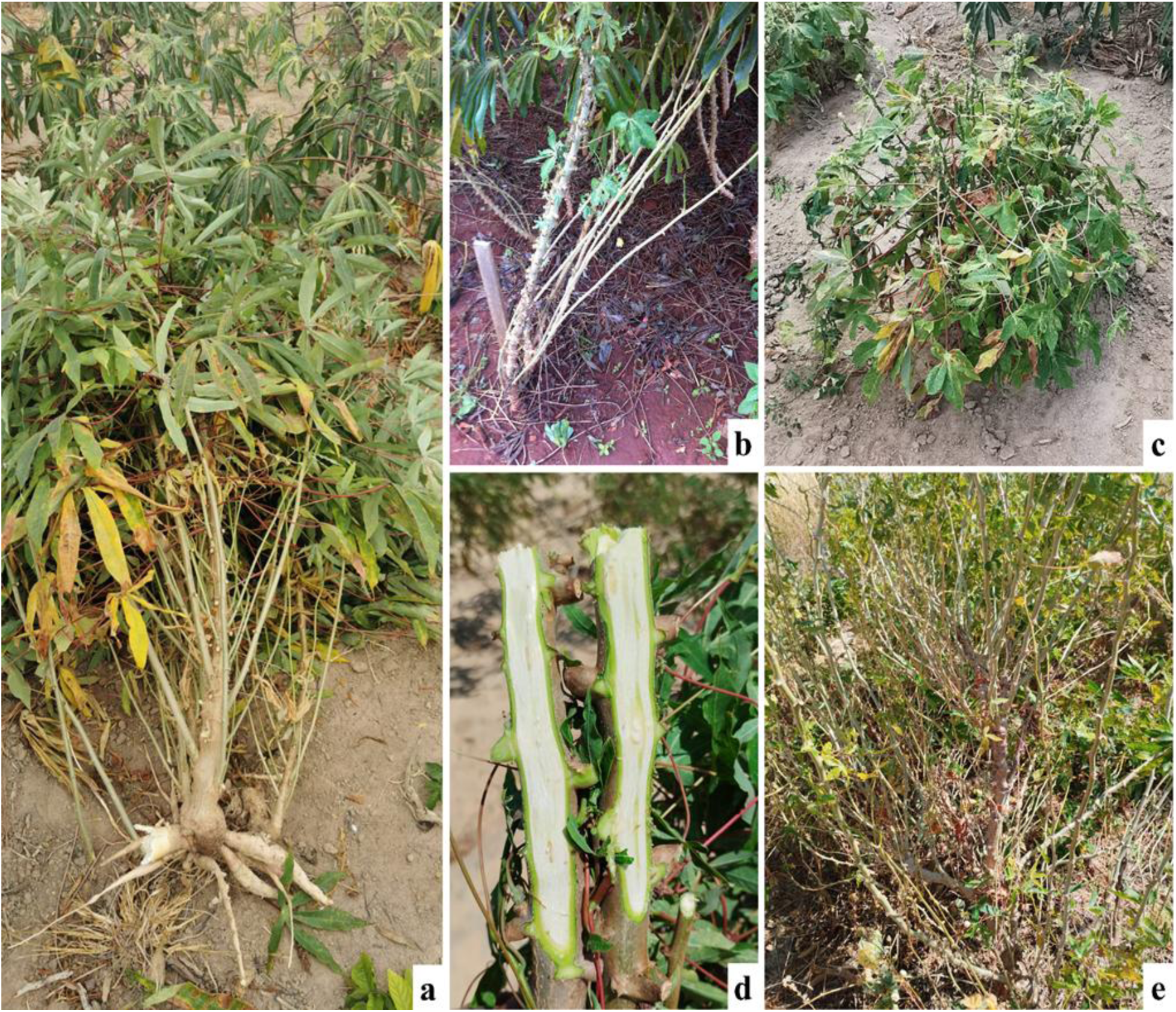
Aspects of cassava plants with symptoms of “Oversprouting”. Plants showing abnormal “bushy” shoot emissions and stunting (dwarfism) of young plants (**c**). Longitudinal section of a cassava stem with a healthy appearance and intact vascular tissue **(d).**

In contrast to “oversprouting”, CWBD, consists of dwarfism and proliferation of weak, spindly sprouts, but the formation of broom-like structures on cassava stems, the short internodes and the striking vascular eventually causing wilting of CWBD affected plants are distinctive features of CWBD.

### Metagenome sequencing for detection of pathogens

Illumina sequencing was conducted on pooled samples from asymptomatic plants, plants exhibiting witches’ brooms (DWBP), and plants showing vascular necrosis (DWBS) (Table 1). The median number of reads obtained per pool was between 48,823,878 and 62,648,108. Approximately 84.91% of the reads from the asymptomatic pool mapped to the cassava genome, compared to 78.78% and 70.76% of the reads from the DWBS and DWBP pools, respectively. *Rhizoctonia*/*Ceratobasidium* contigs were identified exclusively in the DWBP and DWBS pools, with 500 and 495 contigs, respectively (Table 2).

**Table 2.**
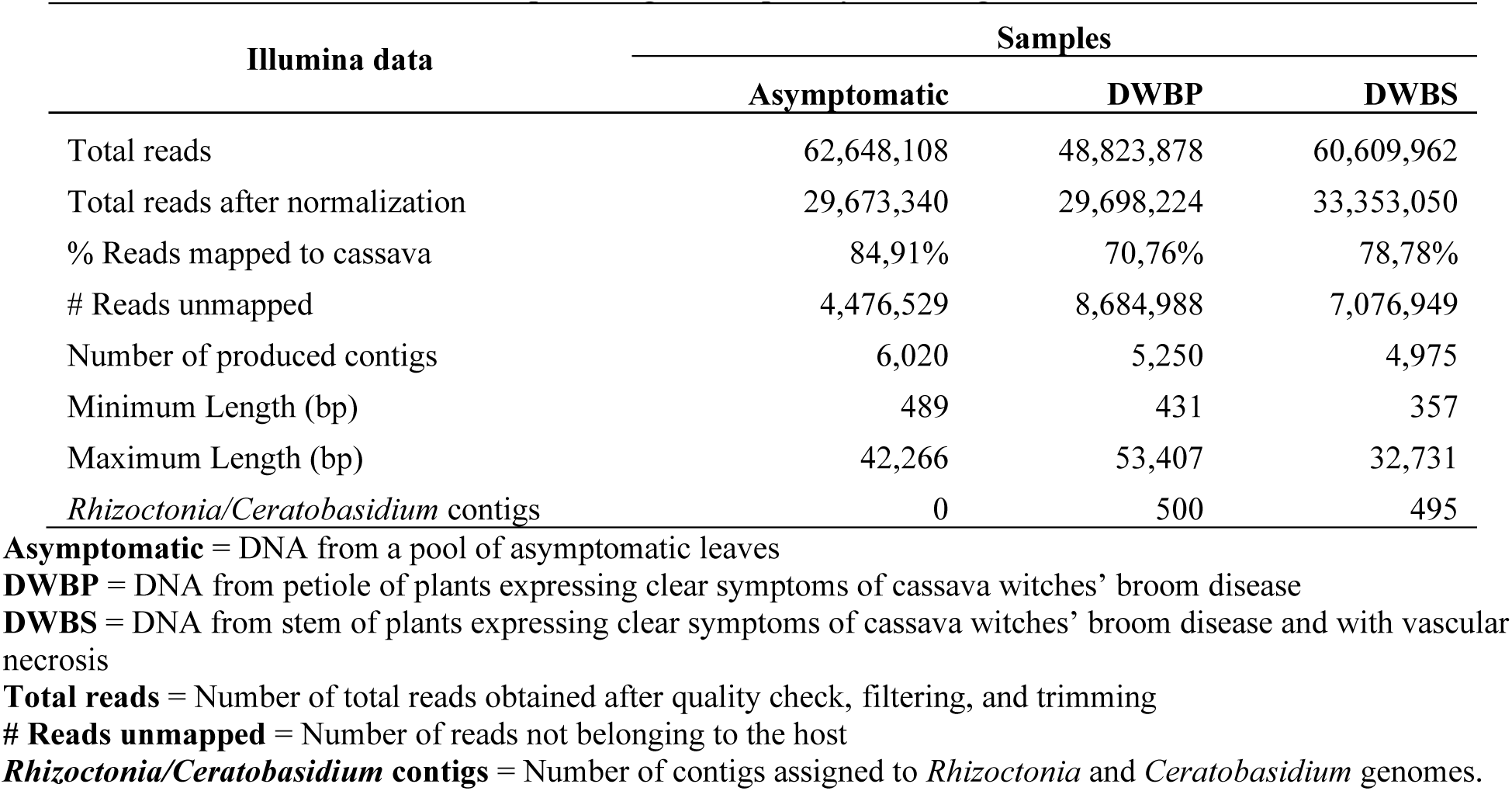
Results from Illumina sequencing after quality filtering.

Microbial community analysis using a metagenome approach was performed to associate organisms with cassava showing witches’ broom symptoms (Figure 4A). In asymptomatic samples and DWBP, fungi of the cosmopolitan Glomerellaceae (genus Ascomycota) were most prevalent, with substantial relative abundance of 87.22 and 47.02%, respectively. In the DWBS pools, 57% of the reads were assigned to fungi of the Ceratobasidiaceae which were also abundant in DWBP (27.15%). Ceratobasidiaceae were exclusively found in the DWBP and DWBS pools, suggesting a causal association with these diseases. Other prominent families, including Glomerellaceae and Botryosphaeriaceae, were also well represented in the asymptomatic and DWBP samples, whereas Diaporthaceae and Hymenochaetaceae appeared less frequently and were specific to certain diseased pools. The distribution of microbial genera is detailed in Figure 4B.

**Figure 4.**
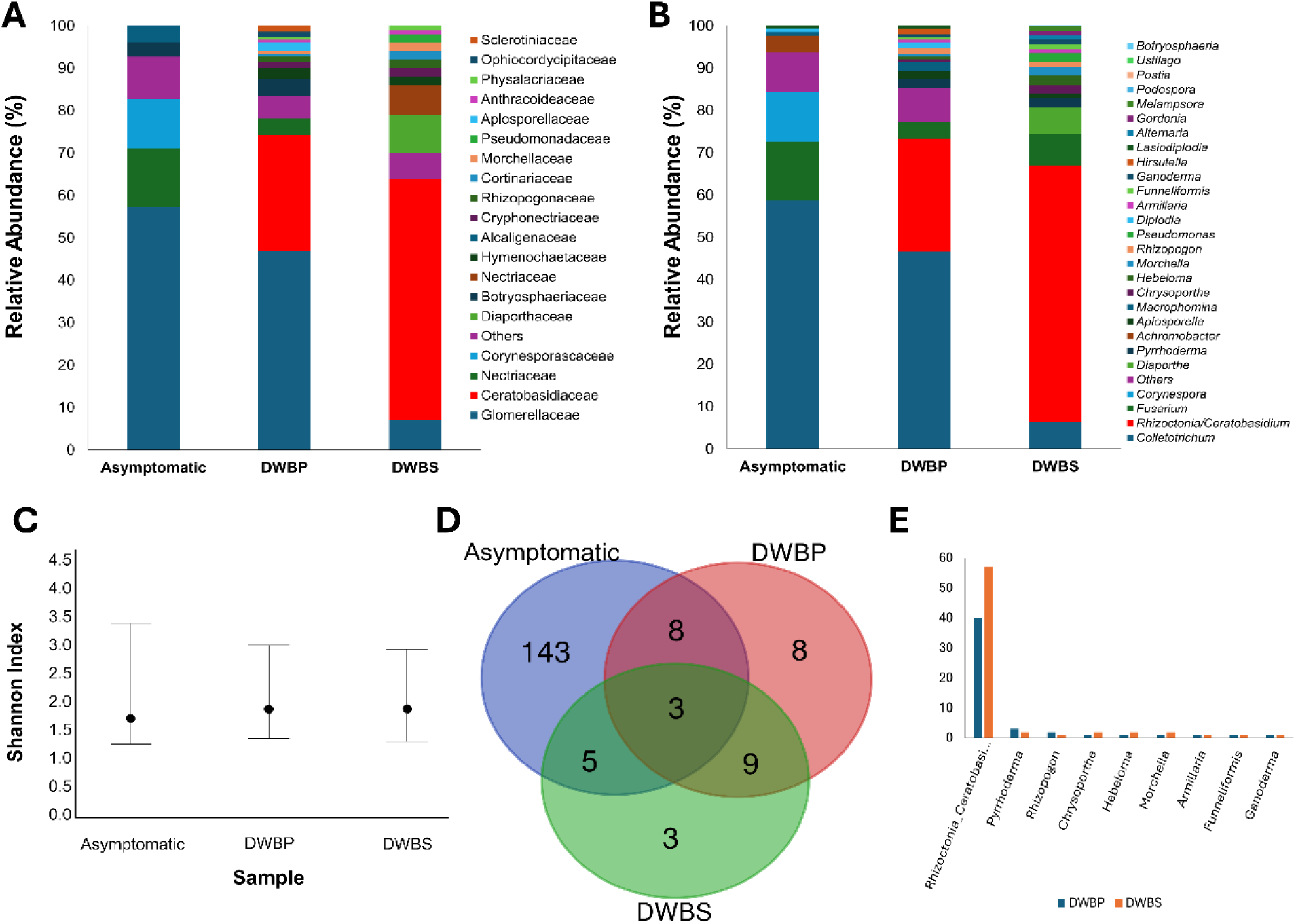
Taxonomic composition of leaves of asymptomatic plants; petioles (DWBP) and stems (DWBS) of plants with symptoms of cassava witches’ broom disease and vascular necrosis. (**A)** Relative abundance at the family level; (**B**) Relative abundance at the genus level; (**C)** Venn diagram with genus distributions among asymptomatic plants and plants expressing clear symptoms of cassava witches’ broom disease and vascular necrosis (numbers inside the diagram indicate the numbers of exclusive or shared OTUs); (**D)** Graphical representation of taxa identified in DWBP and DWBS.

Sequences of fungi from the genus *Colletotrichum* were the most abundant in asymptomatic samples (58.7% of reads) and were also present in DWBP and DWBS (46.67 and 6.38% of reads, respectively). Sequences from several fungal genera, including *Aplosporella*, *Chrysoporthe*, *Hebeloma*, and *Morchella*, were uniquely found in DWBP and DWBS pools (Figure 4 a), suggesting an association with disease conditions.

Sequences from the *Rhizoctonia/Ceratobasidium* genus were the most abundant in the DWBP and DWBS pools, accounting for 26.67% and 60.58% of the reads, respectively, whereas they were absent in asymptomatic sample pools. This distribution suggests a potential relationship between this genus and disease. *Fusarium* was another prevalent genus, found in high abundance in asymptomatic samples (13.87%) and DWBS (7.44%) but was less common in DWBP (4%). Other genera, such as *Corynespora*, *Diaporthe*, and *Pyrrhoderma,* showed specific distributions; *Corynespora* was exclusive to asymptomatic samples, *Diaporthe* was found only in DWBS, and *Pyrrhoderma* was present in DWBP (Figure 3).

Shannon Diversity Index values for microbial families were 1.71 for asymptomatic samples, 1.80 for DWBP, and 1.79 for DWBS (Figure 4C), reflecting similar diversity levels among sample groups with slightly higher diversity in DWBP. Bootstrap confidence intervals ranged from 1.25 to 3.46 for asymptomatic, 1.44 to 3.01 for DWBP, and 1.30 to 2.94 for DWBS, showing no significant differences in microbial diversity among samples.

Graphical representation in Figure 4D illustrates that asymptomatic plants had the highest number of unique taxa (n=143), whereas DWBP and DWBS had fewer unique taxa (n=8 and n=3, respectively). Nine genera were shared exclusively between DWBP and DWBS, suggesting a common pathogen composition in both conditions.

In infected samples from DWBP and DWBS, the genus *Rhizoctonia/Ceratobasidium* was the most abundant, with 38 and 57 contigs matching database entries (Figure 4E), whereas *Pyrrhoderma*, *Chrysoporthe*, *Hebeloma*, *Morchella*, *Rhizopogon*, *Armillaria*, *Funneliformis*, and *Ganoderma* were present in low abundance but absent in asymptomatic samples.

Nanopore sequencing confirmed a high number of reads for *Ceratobasidium/Rhizoctonia* genetic material in DWBP and DWBS, indicating that both conditions had similar or identical pathogens.

### Phylogenetic analyses

Previous research showed that this *Ceratobasidium* genus was associated with witches broom disease. Hence, *Rhizoctonia/Ceratobasidium* sequences from the ITS rDNA region, including partial sequences of the 18S and 28S rDNA genes, the internal transcribed spacers 1 and 2 (ITS1 and ITS2), and the 5.8S rDNA gene were analyzed. Sequences from HTS mapped against a randomly chosen *Rhizoctonia solani* reference sequence (GenBank accession no. AF153800). The resulting consensus sequences were used for subsequent phylogenetic analysis. In the alignment matrices, sequences from the first report of *Ceratobasidium* sp. associated with cassava in Asia (Leiva et al., 2023) were used as references (Table 3).

**Table 3.** Description of the isolates used for phylogenetic analysis based on the ITS rDNA region of *Rhizoctonia-like* species, with collection details and GenBank accession numbers.

| GenBank | Taxonomic assignation | Location | Host | Isolation source |
| --- | --- | --- | --- | --- |
| AB122145 | *Ceratobasidium* sp. AG-k | Japan | *Allium cepa* | mycelium |
| AB196647 | *Ceratobasidium* sp. | Japan | *Arachis hypogaea* | mycelium |
| AB196650 | *Ceratobasidium* sp. AG-I | Japan | *Artemisia* sp. | mycelia |
| AJ427407 | *Ceratobasidium* sp. AGR | USA | *Cucumis* sp. | mycelia |
| AF354095 | *Ceratobasidium* sp. AGQ | Japan | Cynodon_Soil | mycelia |
| KY075643 | *Ceratobasidium theobromae* | China | *Lonicera japonica* | -- |
| KP119764 | *Ceratobasidium theobromae* | China | *Lonicera japonica* | -- |
| KP119766 | *Ceratobasidium theobromae* | China | *Lonicera japonica* | -- |
| KP119765 | *Ceratobasidium theobromae* | China | *Lonicera japonica* | -- |
| OR145521 | *Ceratobasidium theobromae* | Cambodia | *Manihot esculenta* | petiole |
| OR145522 | *Ceratobasidium theobromae* | Vietnam | *Manihot esculenta* | petiole |
| OR145523 | *Ceratobasidium theobromae* | Lao PDR | *Manihot esculenta* | petiole |
| AB286940 | *Ceratobasidium* sp. AG-P | Japan | -- | -- |
| AB290017 | *Ceratobasidium* sp. AG-E | Japan | -- | -- |
| AB000019 | *Rhizoctonia solani* | Japan | -- | -- |
| AY154313 | *Thanatephorus cucumeris* | Brazil | -- | -- |
| AF354108 | *Thanatephorus cucumeris* | Veracruz | *Solanum tuberosum* | mycelia |
| AF153804 | *Rhizoctonia solani* | Australia | *Pterostylis acuminata* | -- |
| AB286930 | *Ceratobasidium* sp. AG-Ba | Japan | Rice | -- |
| AB196659 | *Ceratobasidium* sp. AG-T | Japan | *Rosa odorata* | -- |
| AB196653 | *Ceratobasidium* sp. AG-L | Japan | Soil | mycelium |
| AB000003 | *Rhizoctonia solani* | Japan | Soil | -- |
| AF153800 | *Rhizoctonia solani* | Australia | Soil | -- |
| AB196649 | *Ceratobasidium* sp. AG-H | Japan | Soil | mycelia |
| AB290021 | *Ceratobasidium* sp. AG-C | Japan | Sugar beet | mycelia |
| KU255726 | *Ceratobasidium theobromae* | Indonesia | *Theobroma cacao* | leaf |
| KU319572 | *Ceratobasidium theobromae* | Indonesia | *Theobroma cacao* | leaf |
| KU255724 | *Ceratobasidium theobromae* | Indonesia | *Theobroma cacao* | petiole |
| KM523668 | *Ceratobasidium theobromae* | Indonesia | *Theobroma cacao* | mycelium |
| KU319573 | *Ceratobasidium theobromae* | Indonesia | *Theobroma cacao* | leaf |
| KU310940 | *Ceratobasidium theobromae* | India | *Theobroma cacao* | petiole |
| KU310941 | *Ceratobasidium theobromae* | India | *Theobroma cacao* | petiole |
| AB000004 | *Rhizoctonia solani* | U.S.A. | Tobacco | -- |
| AB198702 | *Ceratobasidium* sp. AG-D | Japan | *Zoysia japonica* | mycelia |
| MZ159527 | *Scotomyces subviolaceus* | Scotland | -- | -- |
| MT340869 | *Heteroacanthella acanthophysa* | -- | -- | -- |
| PQ192611 | *Ceratobasidium theobromae* | Brazil | *Manihot esculenta* | petiole |
| PQ192612 | *Ceratobasidium theobromae* | Brazil | *Manihot esculenta* | stem |

The two ITS rDNA consensus sequences assembled from the DWBS and DWBS samples were placed in a well-supported clade comprising only *C. theobromae* (Figure 5). The subclade formed assigned isolates from cassava and cacao associated with witches’ broom disease (CWB) and vascular streak dieback (VSD), respectively.

**Figure 5.**
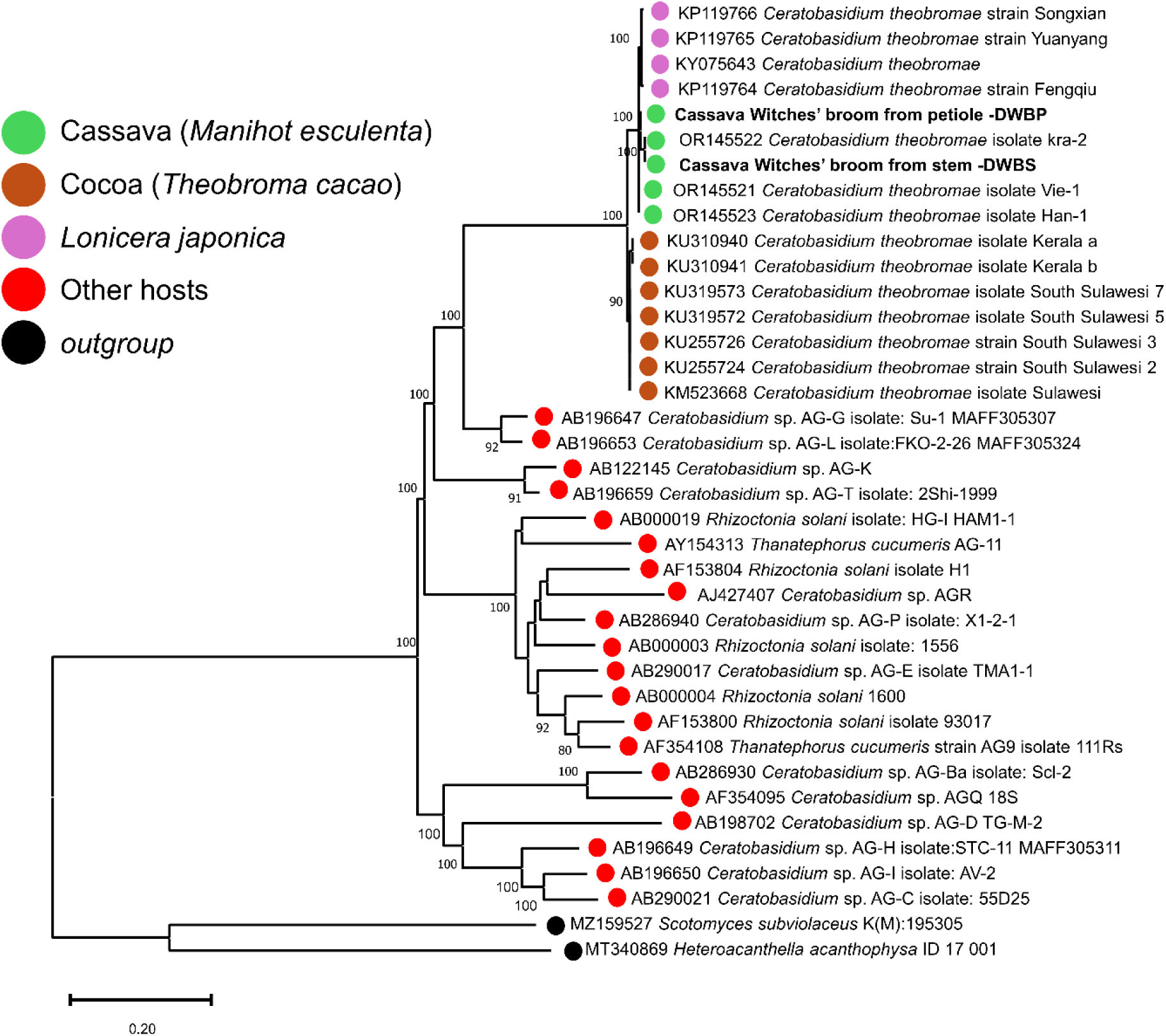
Relationship between *Rhizoctonia/Ceratobasidium* sp. from cassava collected from indigenous villages in the Brazilian Rainforest in the Oiapoque region and others ‘*Rhizoctonia*-like*’* species sequences available in GenBank based on the Internal Transcribed Spacer (ITS) sequence of the rDNA. A set of isolates found causing vascular necrosis in cacao and honeysuckle and witches’ broom in cassava in Asian countries were also included. The tree was constructed using a maximum-likelihood method based on the Tamura 3-parameter with gamma distribution (T92+G) model with 1000 bootstrap replicates. Bootstrap values higher than 70% are displayed close to the branch. *Scotomyces subviolaceus* and *Heteroacanthella acanthophysa* were used as outgroups.

The molecular markers evaluated for *R. theobromae* from Brazil had sequence identities of 95.31% for *atp6*, 96.88% for *tef1*, 97.99% for *rpb2,* and 98.22% for SSU, with the *C. theobromae* LAO1 reference genome (GCA_037974915.1). This provides molecular evidence that the fungus associated with WBD of cassava in Brazil is *R. theobromae* (*C. theobromae*).

### PCR-based detection

To further support the hypothesis that *R. theobromae* (*C. theobromae*) is the cause of this epidemic outbreak in the state of Amapá, Brazil, PCR analyses using specific primers (CWBD-CIAT-F2/R2) described by Leiva et al. (2023) were used. This PCR primer set targeted the coding region of a putative Ca2+/calmodulin-dependent protein kinase gene (CaMK) identified in *C. theobromae* (GenBank accession no. KAB5596398).

Cassava samples collected from indigenous villages affected by the CWBD outbreak, including the leaves, petioles, and stems of symptomatic and asymptomatic plants, were subjected to PCR. To prove specificity, plants displaying symptoms of anthracnose, frogskin disease, and oversprouting were randomly collected from the Embrapa Cassava Germplasm Bank in Cruz das Almas, Bahia, Brazil and evaluated.

From 50 cassava samples with disease symptoms, 37 tested positive for *R. theobromae*, indicating a significant correlation between the pathogen and the disease. Conversely, among the 15 asymptomatic samples, only one tested positive and there were no false-positives among the 40 samples from healthy plants collected from the Germplasm Bank in Cruz das Almas, Brazil (located over 2,500 km away from the affected fields) (Table 4). These results underscore the strong association of *R. theobromae* (*C. theobromae*) with CWBD, particularly in diseased samples from the outbreak region.

**Table 4.** PCR analysis of field samples collected on the indigenous villages that are suffering with the outbreak (leaves, petioles, and stem from plants with and without disease symptoms) showed a strong association of *Rhizoctonia theobromae* (*Ceratobasidium theobromae*) with CWBD. As means of comparison, plants randomly collected from the Embrapa Cassava Germplasm Bank expressing symptoms of anthracnose, frogskin disease and Oversprouting were included.

| Phenotype | Number of samples | PCR (–) | PCR (+) |
| --- | --- | --- | --- |
| Asymptomatic | 15 | 14 | 1 |
| Diseased | 50 | 13 | 37 |
| Healthy* | 40 | 40 | 0 |

\*Plants collected in the Cassava Germplasm Bank in Cruz das Almas, Brazil, more than 2.500 km from the diseased fields, were considered healthy plants for *R. theobromae* infection.

### Rhizoctonia theobromae genetic diversity analysis

The putative pathogenicity gene (CAMK/CMKL) used for species-specific detection of *R. theobromae* was used for population analysis to provide an insight into the genetic diversity of the fungus in Brazil and its relationship with *R. theobromae* (*C. theobromae*) fungi causing cassava witches’ broom disease in Asia. A total of 18 high-quality CAMK/CMKL consensus sequences were retrieved from the samples and used for genetic diversity analysis (Table 5).

**Table 5.** Summary of cassava isolates used in the study of the diversity of putative Ca²⁺/calmodulin-dependent protein kinase genes (CAMK/CAMKL). The table includes information on sample identification, haplotype, cultivar, year of collection, country, specific location, tissue type, and corresponding GenBank accession numbers for isolates from various Asian countries and Brazil.

| Sample | <sup>1</sup>Hap. | Cultivar | Year | Country | Location | Tissue | GenBank |
| --- | --- | --- | --- | --- | --- | --- | --- |
| AP-2204 | H1 | Bâton-Uaçá | 2023 | Brazil | Ariramba | Stem | PQ202677 |
| AP-2207 | H1 | Bâton-Ló | 2023 | Brazil | Ariramba | Stem | PQ202678 |
| AP-2303 | H1 | Tête-Bleu | 2023 | Brazil | Galibi | Stem | PQ202679 |
| AP-2305 | H1 | Tête-Bleu | 2023 | Brazil | Galibi | Stem | PQ202680 |
| AP-2351 | H1 | Bolinha | 2023 | Brazil | Galibi | Petiole | PQ202682 |
| AP-2353 | H1 | Tête-Bleu | 2023 | Brazil | Galibi | Petiole | PQ202683 |
| AP-2602 | H1 | Kawa'wa | 2023 | Brazil | Ahumã | Petiole | PQ202684 |
| AP-2616 | H1 | Kalixá | 2023 | Brazil | Ahumã | Petiole | PQ202685 |
| AP-2617 | H1 | Kawa'wa | 2023 | Brazil | Ahumã | Stem | PQ202686 |
| AP-2663 | H1 | Camarão | 2023 | Brazil | Ahumã | Stem | PQ202687 |
| AP-2706 | H1 | Tucumã | 2023 | Brazil | Anawerá | Leaf | PQ202688 |
| AP-2711 | H1 | Tucumã | 2023 | Brazil | Anawerá | Stem | PQ202689 |
| AP-2713 | H1 | Xingú | 2023 | Brazil | Anawerá | Leaf | PQ202690 |
| AP-2719 | H1 | Xingú | 2023 | Brazil | Anawerá | Stem | PQ202691 |
| AP-2802 | H1 | Manteiga | 2023 | Brazil | Yanawaka | Stem | PQ202692 |
| AP-2807 | H1 | Manteiga | 2023 | Brazil | Yanawaka | Petiole | PQ202693 |
| AP-2809 | H1 | Pretinha | 2023 | Brazil | Yanawaka | Petiole | PQ202694 |
| AP-2812 | H1 | Manteiga | 2023 | Brazil | Yanawaka | Stem | PQ202695 |
| YBa-1 | H3 | KM 94 | 2012 | Vietnam | Yen Bai | Stem | OQ863061 |
| YBa-2 | H4 | KM 94 | 2012 | Vietnam | Yen Bai | Stem | OQ863062 |
| YBa-3 | H6 | KM 94 | 2012 | Vietnam | Yen Bai | Stem | OQ863063 |
| YBa-4 | H1 | KM 94 | 2012 | Vietnam | Yen Bai | Stem | OQ863064 |
| YBa-5 | H2 | KM 94 | 2012 | Vietnam | Yen Bai | Stem | OQ863065 |
| HBi-1 | H3 | SC 205 | 2012 | Vietnam | Hoa Binh | petiole | OQ863066 |
| HBi-2 | H3 | SC 205 | 2012 | Vietnam | Hoa Binh | petiole | OQ863067 |
| DNa-1 | H3 | KM 94 | 2012 | Vietnam | Dong Nai | Stem | OQ863068 |
| DNa-2 | H3 | KM 94 | 2012 | Vietnam | Dong Nai | petiole | OQ863069 |
| DNa-3 | H1 | KM 94 | 2012 | Vietnam | Dong Nai | Stem | OQ863070 |
| Kra-1 | H1 | NA | 2013 | Cambodia | Kratie | Root | OQ863071 |
| KCh-1 | H3 | NA | 2013 | Cambodia | Kampong Chan | Root | OQ863072 |
| PVe-1 | H3 | NA | 2013 | Cambodia | Prey Veng | Root | OQ863073 |
| TNi-1 | H3 | NA | 2013 | Vietnam | Tay Ninh | Root | OQ863074 |
| TNi-2 | H9 | NA | 2013 | Vietnam | Tay Ninh | Root | OQ863075 |
| Min-1 | H9 | NA | 2017 | Philippines | Mindanao | NA | OQ863076 |
| Min-2 | H8 | NA | 2017 | Philippines | Mindanao | NA | OQ863077 |
| Min-3 | H10 | NA | 2017 | Philippines | Mindanao | NA | OQ863078 |
| Ray-1 | H10 | HB 60 | 2017 | Thailand | Rayong | Root | OQ863079 |
| Ray-2 | H10 | HB80 | 2017 | Thailand | Rayong | Root | OQ863080 |
| Ray-3 | H10 | <sup>2</sup>Ray.7 | 2017 | Thailand | Rayong | Root | OQ863081 |
| Ray-4 | H8 | Ray.86-13 | 2017 | Thailand | Rayong | Root | OQ863082 |
| Sal-1 | H1 | NA | 2020 | Lao PDR | Salavan | Leaf | OQ863083 |
| Sal-2 | H3 | NA | 2020 | Lao PDR | Salavan | Leaf | OQ863084 |
| Vie-1 | H7 | NA | 2022 | Lao PDR | Vientiane | Stem | OQ863085 |
| Vie-3 | H1 | KU50 | 2023 | Lao PDR | Vientiane | Stem | OQ863086 |
| Vie-4 | H5 | Ray.11 | 2023 | Lao PDR | Vientiane | Stem | OQ863087 |

<sup>1</sup>**Hap.** = Haplotype; <sup>2</sup>**Ray.** = Rayong

Genetic analysis of the *R. theobromae* populations revealed a complex population structure with notable variations in genetic diversity across different regions. In Brazil, the fungal isolates exhibited very low genetic diversity, with a Shannon diversity index of 0.12 and a nucleotide diversity (π) of 0.001, suggesting minimal variation within this region. The haplotype diversity in Brazil was 0.0, indicating a single haplotype, while the Philippine isolates displayed significantly higher genetic diversity, with a Shannon index of 1.56, nucleotide diversity of 0.025, and haplotype diversity of 0.85.

Significant genetic differentiation between populations, indicated by an F_ST_ value of 0.68, underscores the potential influence of geographical and environmental factors on the evolution and spread of *R. theobromae*. The low gene flow between populations, with an estimated migration rate (Nm) of 0.5, suggests relative isolation of these groups.

The high concentration of individuals in the H1 haplotype, particularly in Brazil, may indicate a common ancestor or founder population (Figure 6A). The H1 haplotype is shared by Vietnam, Cambodia, and Laos, suggesting genetic connections or common ancestor. This central haplotype is connected to other less frequent haplotypes, such as H2, H3, H4, and H5, which are found in more restricted populations.

**Figure 6.**
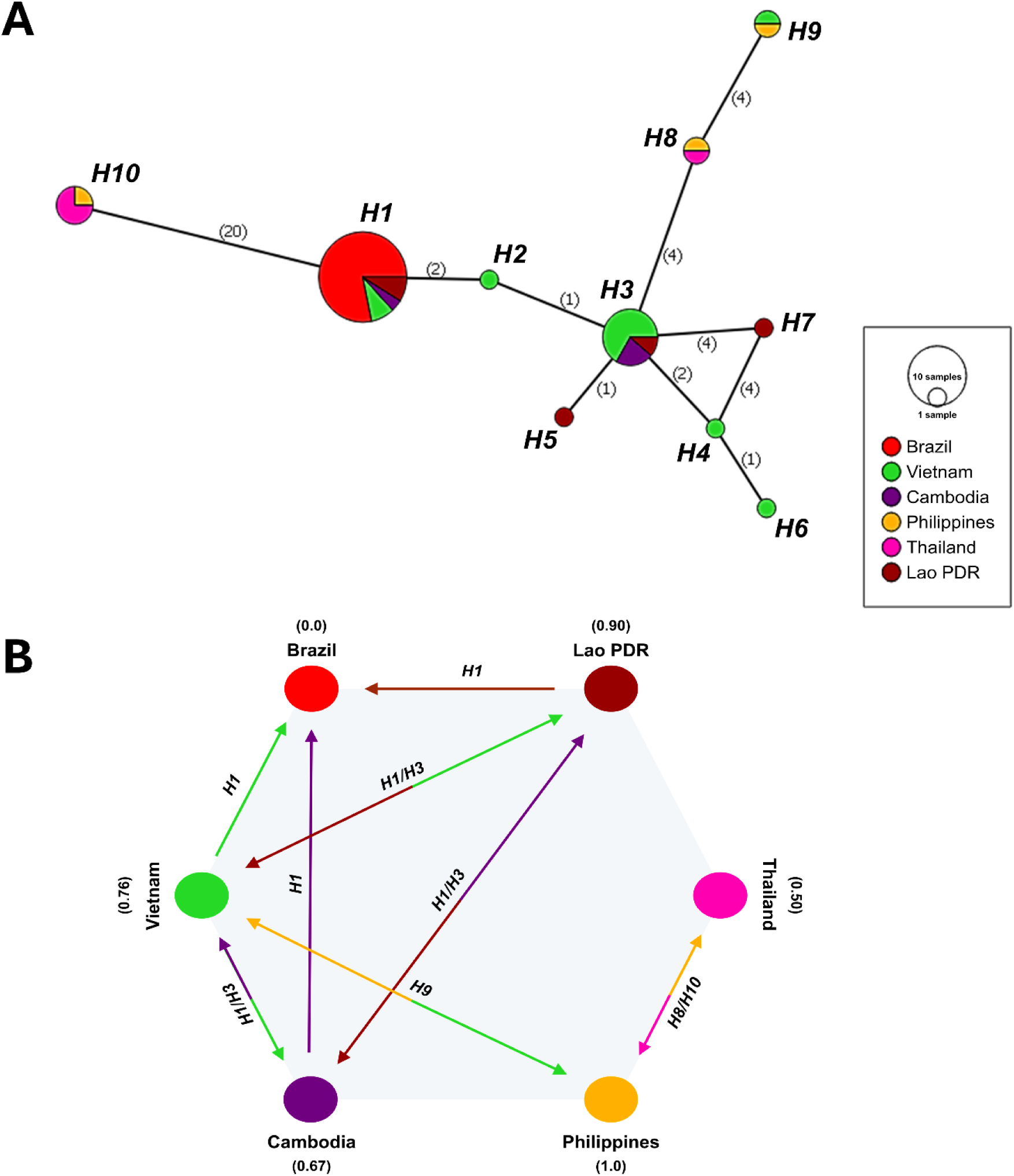
**A.** Haplotype network of *Rhizoctonia theobromae* isolates based putative Ca^2+^/calmodulin-dependent protein kinase gene (CAMK/CAMKL). The network shows the number of individuals for each haplotype (circle size) and population of origin (color). Numbers in parentheses next to the branches indicate the number of mutations that separated the different haplotypes. **B.** Diagram illustrating the sharing of haplotypes between the five Asian populations and Brazil. Values in parentheses next to the names indicate haplotype diversity (Hd) estimated for the population.

The H10 haplotype, which is distant from the others, suggests greater evolutionary divergence or an older founding event. Haplotype diversity was highest in the Philippines and, to a lesser extent, in Brazil and Laos, while Thailand showed the lowest diversity. The H3 haplotype is shared between Vietnam and Cambodia, indicating genetic proximity. The connection between Thailand and the Philippines for the H8/H10 haplotypes suggests genetic sharing, possibly due to spore dispersal and/or transit of infected plants (Figure 6B).

## DISCUSSION

The new cassava disease found in indigenous villages in Oiapoque, Amapá, Brazil shows all characteristics of Cassava Witches’ Broom Disease (CWBD). The disease manifests as stunted growth, proliferation of weak, thin shoots, formation of broom-like structures on cassava stems, followed by chlorosis, wilting, leaf drying, and apical death, leading to the most severe impact on cassava cultivation.

In Brazil, cassava plants affected by witches’ broom disease display, as previously described (Silberschmidt and Campos 1944; Flores et al. 2013), and share some symptoms with CWBD, such as leaf chlorosis, abnormal shoot proliferation, and stunted growth. However, the crucial distinction is the absence of the ‘broom-like’ leaf aggregation (rosette) and vascular necrosis. Symptoms can vary with the environment and genotype; however, molecular evidence has shown a distinct phytoplasma (16SrIII-B) associated with oversprouting disease (Flores et al. 2013) and *Ceratobasidium theobromae* (*Rhizoctonia theobromae*) associated with CWBD (Leiva et al. 2023; Gil-Ordóñez et al. 2024).

The metagenome analysis of the microbial communities associated with witches’ broom disease of cassava in Brazil revealed a similar composition of the microbiota, although specific genera such as *Rhizoctonia/Ceratobasidium* were notably more abundant in symptomatic samples.

The identification of several genera, including *Rhizoctonia*/*Ceratobasidium*, *Aplosporella*, *Chrysoporthe*, and *Hebeloma*, only found in diseased samples indicates that specific microbial communities are associated with disease status, while *Rhizoctonia*/*Ceratobasidium* was significantly correlated with CWBD symptoms. *Rhizoctonia theobromae* was identified as a novel cassava pathogen in Asia (Leiva et al. 2023) and, more recently, in French Guiana (Pardo et al. 2024), marking a significant finding for this region.

Phylogenetic analysis further supported the association between *R. theobromae and* CWBD, and clustered *R. theobromae* from cassava in Brazil, along with other isolates from cassava and cacao in Asia. The ITS phylogeny contributes significantly to the understanding of the diversity and association of Rhizoctonia species by providing a molecular framework for delineating and classifying the taxa within this group. Gónzalez et al. (2016) demonstrates the utility of ITS sequences, among other loci, in resolving phylogenetic relationships within the *Rhizoctonia* complex. This confirmed the monophyly of the family *Ceratobasidiaceae* and revealed well-supported monophyletic groups consistent with the anastomosis groupings.

*Ceratobasidium theobromae* is the causative pathogen for cassava witches’ broom disease (CWBD) and vascular streak dieback (VSD) of cocoa (Guest and Keane 2018; Leiva et al. 2023; Gil-Ordóñez et al. 2024; Pardo et al. 2024). A paper published by Oberwinkler et al. (2013) highlighted the complex taxonomy of this species, with *Ceratobasidium* representing the teleomorphic (sexual) state, whereas the anamorphic (asexual) state is classified as *Rhizoctonia*. Additionally, some studies have grouped similar fungal lineages, including *C. theobromae*, within *Thanatephorus*, which also represents a teleomorphic genus closely related to *Ceratobasidium*. This overlapping classification underscores the importance of precise molecular and phylogenetic analyses to resolve taxonomic ambiguities. For consistency, the use of *Rhizoctonia theobromae* aligns with current conventions for classifying the anamorphic stage of the pathogen, and was therefore used as the preferred name in this study.

PCR analysis confirmed the role of *R. theobromae* in CWBD. A strong association was observed between the incidence of the pathogen and the disease, with a high prevalence in cassava samples from the affected villages. This result was consistent with the observed symptomology and sequencing data, affirming that *R. theobromae* is a key pathogen in CWBD.

Analysis of the genetic diversity of the Ca²⁺/calmodulin-dependent protein kinase (CaMK) gene showed a high degree of identity between Brazilian and Asian isolates, indicating a common genetic origin. This information is crucial for understanding the global distribution of the pathogen and its genetic variability and can inform management strategies and further research.

The integration of symptom observation, high-throughput sequencing, and PCR-based detection has elucidated the role of *C. theobromae* in the cassava witches’ broom disease in the Oiapoque region of Amapá, Brazil. This is the first report of the presence of *C. theobromae* in Brazil, which represents an imminent risk to the cassava crop. This pathogen can spread through infected plant materials, pruning tools, infested soil, and probably by the wind (thought basidiospores). Most critically, the movement of the propagating material between regions can facilitate the spread of pathogens. Mitigation measures must be adopted, comprising an intensified monitoring of cultivation areas, quarantine measures to restrict the movement of plant materials, the use of healthy cuttings produced in disease-free regions, application of specific fungicides, removal and burning of diseased plants, asepsis of tools, and alerting the local population and transiting people.

*Rhizoctonia theobromae,* (syn = *Ceratobasidium theobromae* or *Onobasidium theobromae* or *Thanatephorus theobromae*) has been classified as a quarantine pest by the Brazilian Ministry of Agriculture and Livestock (MAPA), a "Technical Communication Note" was sent to the MAPA-registered SEI: 21157.001205/2024-84, as documented in the files 10729769, 10730668, and 10730758. The presence of this pathogen was confirmed by an independent laboratory registered under the numbers N° 03575/24-GO and N° 03575/24-GO (supplementary file), and quarantine measures were implemented (MAPA, 2024). The presence of *R. theobromae* in the Amapá region represents a serious threat to cassava farming but also to cultivation of other crops of great economic importance for the country, such as cocoa (*Theobroma cacao*) and putatively other plants from the same family, such as “cupuaçú” (*T. grandiflorum*). Further research is necessary to clarify whether the cassava pathogen is host-specific or can also be a source of infection for other crops, such as cacao.

While global concerns about cassava pathogens have mainly focused on viral diseases, such as Cassava Brown Streak Disease and Cassava Mosaic Disease (Uke et al. 2022; Hareesh et al. 2023), these findings raise the urgency for tight phytosanitary measures to prevent the spread of emerging threats across continents.

## CONCLUSION

Cassava Witches’ Broom Disease (CWBD) in the Oiapoque region of Brazil is characterized by stunted growth, abnormal shoot proliferation, and distinct broom-like structures, with vascular necrosis associated with C. theobromae considered the causal agent of the disease. Metagenomic and phylogenetic analyses confirmed the association of *Rhizoctonia theobromae* (*Ceratobasidium theobromae*) with CWBD in Brazil, consistent with findings from similar cassava diseases in Southeast Asia and suggesting the introduction of this pathogen from this region. This is further corroborated by the low genetic variability among *R. theobromae* (*C. theobromae*) isolates from Brazil, indicating a recent introduction with a founder effect, in contrast to the high diversity of the pathogen observed in Asia.

## ACKNOWLEDGMENTS

To Empresa Brasileira de Pesquisa Agropecuária (Embrapa), Deutsche Sammlung von Mikroorganismen und Zellkulturen (DSMZ) and Alexander von Humboldt Stiftung (AvH) for funding; to Conselho Nacional de Desenvolvimento Científico e Tecnológico-CNPq (Saulo A.S.O. research productivity fellowship grant n° 315172/2021-5); and to Coordenação de Aperfeiçoamento de Pessoal de Nível Superior-CAPES and Alexander von Humboldt Stiftung (Saulo A.S.O. CAPES/von Humboldt experienced research fellowship grant n° 88881.699110/2022-01).

## AUTHOR CONTRIBUTIONS STATEMENT

S.A.S.O., A.L.L., J.A.S., H.S.R., H.F.S. and C.R.J. were responsible for sample collection and on-site disease identification. S.A.S.O., S.S. and S.W. conceptualized the high-throughput sequencing approach for pathogen identification. Data analysis was carried out by S.A.S.O., P.M. and S.S. while data curation was performed by S.A.S.O., S.S., P.M. and S.W. The methodology was developed by S.W., P.M. and S.S. Funding acquisition was managed by S.A.S.O., C.R.J. and S.W. All authors contributed to the original draft and participated in reviewing and editing the manuscript. S.A.S.O. and S.S. contributed equally to this work.

## DATA AVAILABILITY STATEMENT

The datasets generated and/or analyzed during the current study are available from the corresponding author upon reasonable request.

